# Behavioural flexibility masks delayed costs of environmental instability

**DOI:** 10.64898/2026.08.04.742804

**Authors:** Natacha Rossi, Elizabeth Nicholls

## Abstract

Environmental predictability influences the value of information acquired through experience, yet relatively little is known about how instability in resource characteristics influences behavioural organisation during foraging. We tested whether repeated changes in floral orientation, a manipulation of environmental predictability, affect pollen foraging in bumblebees (*Bombus terrestris*) by exposing naïve workers to either stable floral conditions (single flower orientation) or repeated inter-trial changes in flower orientation (three alternating flower orientations), under constant resource availability. We quantified pollen collection, foraging efficiency, revisitation behaviour, floral coverage and sonication behaviour across three successive foraging trials of either constant or variable flower orientation, before assessing performance in a common post-treatment preference test in which bees were offered all three flower orientations and higher pollen rewards. Environmental instability altered the organisation of foraging behaviour. Bees exposed to unstable floral conditions progressively reduced flower revisitation behaviour and were less likely to perform sonication, although floral coverage, defined as the number of unique flowers visited, remained unchanged. Contrary to our predictions, instability had only weak immediate effects on pollen acquisition and foraging efficiency compared to bees tested under stable conditions. However, previous exposure to instability generated carry-over effects in the common post-treatment preference test. Bees previously exposed to unstable conditions were significantly less likely to return with pollen and consequently collected less pollen overall than bees previously exposed to stable conditions. Our results demonstrate that environmental instability can influence pollen foraging in ways that are not captured by immediate measures of performance. Although behavioural adjustments appeared to buffer short-term consequences during repeated foraging trials, carry-over effects were evident when bees were later tested in a common high-reward, multi-orientation floral array. These findings highlight the importance of considering both behavioural flexibility and carry-over effects when evaluating how organisms respond to changing environments.

## Introduction

Animals rely on information acquired through experience to exploit resources efficiently, yet the value of that information depends fundamentally on environmental predictability. When environmental conditions remain stable, previous experience can improve future decisions and enhance foraging performance. In contrast, when environments become unstable through repeated change, information acquired through prior encounters may become less reliable, reducing the benefits of relying on learned behavioural routines. Consequently, theoretical and empirical studies increasingly recognise environmental predictability as a key determinant of information use, learning and foraging decisions (Dall et al., 2005; McNamara & Dall, 2010; McNamara & Houston, 1987; Stephens, 1987, 1989, 1991). Despite this growing interest, relatively little is known about how instability in the physical characteristics of resources influences behavioural organisation during foraging.

Pollinator foraging provides an especially useful system in which to investigate these questions. Pollinators depend heavily on information acquired through previous interactions with flowers, including information about floral identity, reward availability, spatial location and handling requirements (Chittka et al., 1999; Chittka & Thomson, 2001; Raine & Chittka, 2007a). Repeated exploitation of profitable floral resources can generate flower constancy and route-based foraging behaviours that improve foraging efficiency by reducing search, travel and handling costs (Chittka et al., 1999; Grüter, 2026; Saleh & Chittka, 2007; Waser, 1986). These benefits, however, depend on the extent to which floral conditions remain predictable through time. Changes in resource characteristics may reduce the future value of previously acquired information and alter how pollinators organise foraging behaviour.

Most studies of pollinator learning and information use have focused on variation among flower species, reward schedules or spatial distributions of floral resources (Chittka & Thomson, 2001; Dunlap et al., 2017; Lihoreau et al., 2012; Raine & Chittka, 2007b). However, pollinators also encounter substantial variation within plant species. Individual flowers can differ in reward state, morphology and physical presentation, including floral orientation and the arrangement of reproductive structures (Armbruster & Muchhala, 2020; Nicholls & Hempel de Ibarra, 2017; Sponsler et al., 2023). Such variation may be particularly important when pollinators collect pollen rather than nectar. Unlike nectar foraging, pollen collection often requires specialised handling and extraction behaviours that depend strongly on interactions between pollinators and floral structures (Nicholls & Hempel de Ibarra, 2017; Vallejo-Marín, 2022). Consequently, changes in floral presentation have the potential to influence both resource acquisition and the usefulness of information gained during previous floral encounters.

The importance of floral presentation is especially evident in buzz-pollinated systems. Many flowering plants restrict pollen release through poricidal anthers that require bees to generate vibrations, a behaviour known as floral sonication or buzz pollination (Vallejo-Marín, 2022). Although sonication behaviour is largely innate, experience can refine how bees interact with flowers during pollen extraction (Morgan et al., 2016). Recent biomechanical work further demonstrates that successful pollen extraction depends strongly on the mechanical coupling between bees and flowers during handling and buzzing behaviours (Woodrow et al., 2024). These findings suggest that changes in floral presentation may influence pollen-foraging performance through their effects on both handling behaviour and pollen extraction.

Floral orientation is one component of floral presentation known to influence pollinator behaviour and plant reproductive success (Armbruster & Muchhala, 2020; Jirgal & Ohashi, 2023; Makino & Thomson, 2012; Nakata et al., 2022; Nevard & Vallejo-Marín, 2022). Experimental studies show that flower orientation can affect pollinator attraction, landing behaviour, handling stability and pollen transfer (Jirgal & Ohashi, 2023; Makino & Thomson, 2012; Nevard C Vallejo-Marín, 2022). Floral orientation can also influence pollination accuracy, pollen export and protection of reproductive structures from environmental stress (Armbruster & Muchhala, 2020; Nakata et al., 2022; Nevard & Vallejo-Marín, 2022).Previous studies have largely treated floral orientation as a fixed trait, however, in natural environments, floral orientation may vary through time as a consequence of plant development, floral movement, weather conditions or mechanical disturbance (Armbruster & Muchhala, 2020; Ruan & da Silva, 2011; Van Der Kooi et al., 2019; Van Doorn & Van Meeteren, 2003). From the perspective of a foraging pollinator, such changes may create instability in floral handling conditions while leaving resource identity and availability unchanged. Floral orientation therefore provides a tractable means of manipulating environmental instability independently of reward quantity or spatial resource distribution, thereby altering the predictability of floral conditions.

Here, we investigated how instability in floral orientation between successive foraging trials influences pollen foraging in naïve *Bombus terrestris* workers visiting artificial flowers that elicited sonication behaviour. Bees were assigned to one of two treatments: stable conditions, in which flower orientation remained constant across successive foraging trials, with individual bees exposed to a single flower orientation, or unstable conditions, in which flower orientation was repeatedly switched between trials, with individual bees exposed to three flower orientations (upwards, downwards and sideways) in total. In both treatments, the spatial position of flowers and the reward quantity remained unchanged. We quantified pollen collection, trial duration, revisitation behaviour, floral coverage and sonication behaviour to determine how instability influenced both foraging performance and behavioural organisation.

Because stable environments increase the value of previously acquired information, whereas changing environments reduce its reliability (Dall et al., 2005; McNamara & Houston, 1987; Stephens, 1987, 1989, 1991), we predicted that repeated inter-trial switching in floral orientation would reduce pollen-foraging efficiency relative to stable floral conditions. By separating multiple behavioural components of pollen foraging, we additionally asked whether environmental instability alters the organisation of foraging behaviour independently of overall pollen acquisition. Finally, because successful pollen extraction depends strongly on how bees mechanically interact with flowers during sonication, we predicted that environmental instability in floral orientation would alter the probability of sonication during foraging (Morgan et al., 2016; Vallejo-Marín, 2022; Woodrow et al., 2024).

## Materials and Methods

### Study animals and housing

Experiments were conducted using naïve colonies of *B. terrestris* (Koppert Biological Systems) supplied by PG Horticulture (James Avery, 5 Brunel Close, Drayton Fields Industrial Estate, Daventry, Northamptonshire NN11 8RB, UK). Colonies were housed in their original commercial boxes and connected via a Perspex corridor (26.5 x 4.2 x 4.2 cm) to a Perspex foraging arena (40 x 35 x 40 cm), which allowed bees to forage for pollen and nectar in a controlled environment. Bees were provided *ad libitum* access to sugar solution (Biogluc, Biobest) and hand-collected cherry pollen (Firman, USA) in the foraging arena. Sugar solution was presented in (10 ml) plastic syringes suspended from the ceiling of the foraging arena with fishing wire and pollen was presented on chenille stems to mimic anthers, which were placed inside plastic drinking cups. Pollen-foraging workers, identified by visible pollen loads, were individually marked with numbered tags for identification (Thorne, UK).

### Experimental setup

Experiments were conducted in a flight cage (72 × 72 × 72 cm) which was connected to the Perspex foraging arena described above via a Perspex corridor (55 x 13.5 x 14 cm) which was placed on the opposite side of the foraging arena to the colony. A system of gates ensured that only one bee entered the flight cage at a time, preventing the use of social information and allowing accurate individual-level measurements. The flight cage was illuminated from above using a 1200 × 600 mm LED panel (4000 K; LED Panel Store, UK).

Artificial flowers were arranged on a vertical transparent Perspex panel facing the entrance to the flight cage. The cage floor and surfaces were cleaned with ethanol between trials to remove potential scent marks and pollen residues. Arena walls were covered with high-contrast Julesz patterns to facilitate spatial orientation (Julesz, 1962). Bee behaviour during foraging trials was recorded from a lateral perspective using a Blackfly S BFS-U3-13Y3C-C USB3 colour camera (FLIR Systems, USA) fitted with a 10–22 mm f/3.5–4.5 lens (Canon Inc., Japan) mounted via an EF-to-C-mount adapter.

### Artificial flowers

Artificial flowers consisted of a 3D-printed, radially symmetrical blue corolla supporting a yellow chenille-stem anther on which a fixed mass of pollen was presented (Fig. 1). The flower shape was inspired by *Prunus* blossoms, which naturally exhibit a range of floral orientations. The flower colour was selected based on the innate preference of naive bumblebees for blue hues (Raine et al., 2006), with the aim of encouraging pollen foraging behaviour. Each anther consisted of a 1.5 cm chenille stem, which held pollen securely while allowing controlled loading and removal (Russell & Papaj, 2016). Flowers were mounted on adjustable ball-head supports (SmallRig, China), allowing them to be oriented upward, downward, or sideways.

**Figure 1.**
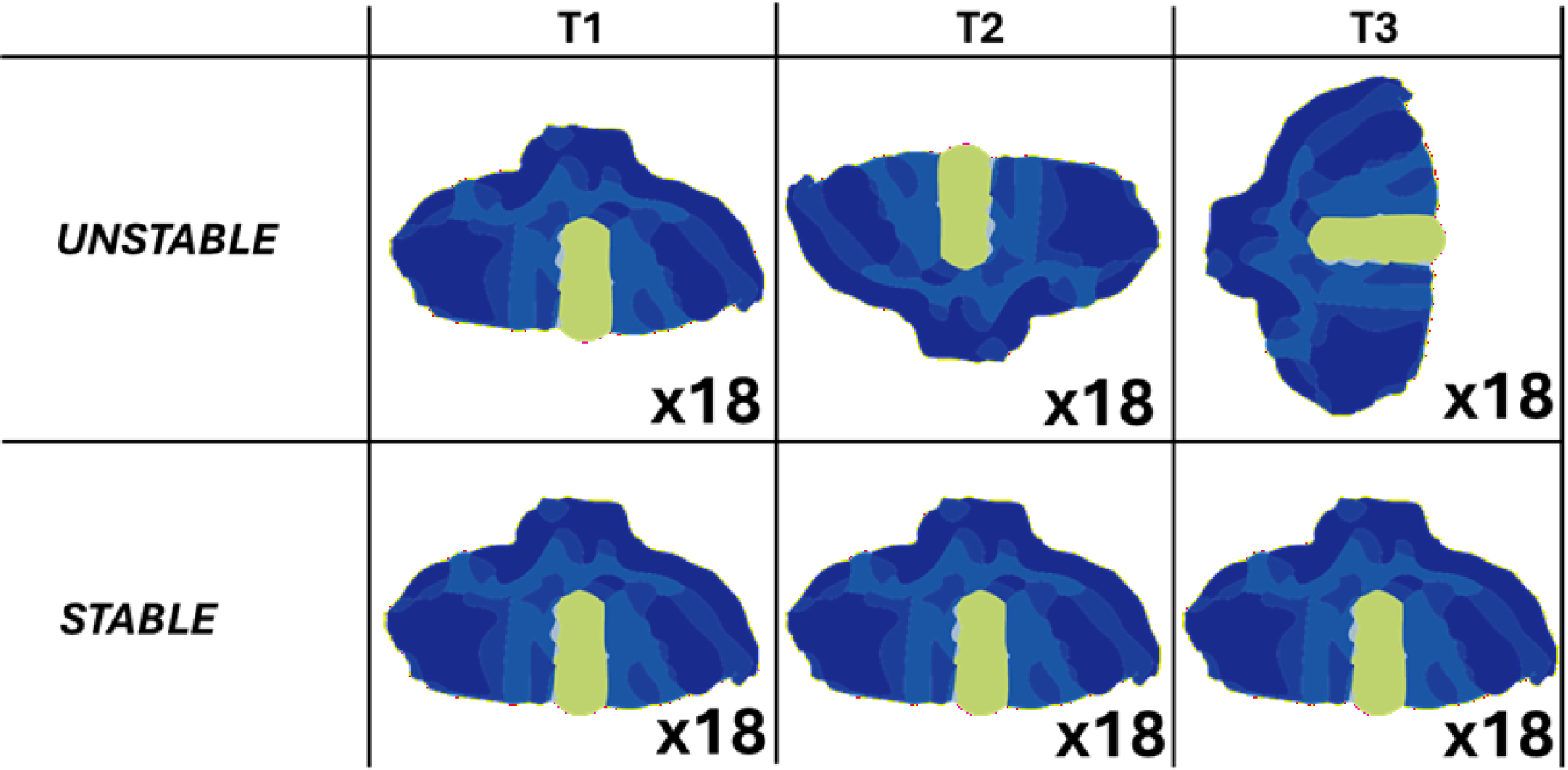
Schematic representation of the experimental treatments across three consecutive foraging trials (T1– T3). Each panel illustrates the orientation of artificial flowers presented within a trial (18 flowers per trial). Flowers consisted of a blue corolla with a central yellow anther (chenille stem loaded with pollen). In the *stable* treatment (bottom row), flower orientation remained the same across all trials. In the *unstable* treatment (top row), flower orientation changed between successive trials.

### Pre-training and orientation preference tests

Workers were pre-trained to collect pollen from a dish placed at the centre of the circular array later used to position artificial flowers, thereby familiarising them with the foraging location prior to testing. Bees that returned subsequently underwent a baseline orientation preference test (P1), followed by three experimental foraging trials (T1–T3) and a post-trial preference test (P2). During preference tests, bees were presented with 12 artificial flowers arranged in a circular array, comprising equal numbers of upward-, downward-, and sideways-facing flowers (n = 4 per orientation). Flowers were positioned in a pseudo-random configuration, and each was loaded with 20 mg of hand-collected cherry pollen, providing pollen in excess of expected foraging demand, based on pilot studies where we calculated that the maximum pollen load weight of a forager was 54 mg. The spatial arrangement of flowers was held constant between the two preference tests.

### Experimental design

Following baseline preference testing, bees were randomly assigned to one of two treatments: (1) a stable treatment, in which bees foraged on flowers of the same orientation across three trials (upward, downward or sideways), or (2) an unstable treatment, in which bees foraged on flowers of three different orientations across successive trials (inter-trial switch).

Before each foraging trial, 18 artificial flowers were arranged in a pseudo-random configuration on two concentric circles (12 + 6 flowers). Compared with the initial and final preference tests, a larger number of flowers was used in foraging trials to encourage bees to forage across many flowers while maintaining realistic pollen rewards per flower. Each flower contained 3 mg of hand-collected cherry pollen, giving a total of 54 mg across the array. As before, this quantity was selected based on pilot observations and designed to encourage bees to visit many flowers within the array. The spatial location of the 18 flowers was held constant across trials; only flower orientation varied according to treatment.

Within the stable treatment, an individual bee was assigned to one of three flower orientations (upward, downward, or sideways) (Fig. 1) and received three trials (T1-3) where flowers remained in this same orientation throughout (e.g. three trials with upward flowers). In the unstable treatment, bees received three trials (T1–T3) in which flower orientation changed between trials (e.g. Trial 1 = upwards, Trial 2 = downward, Trial 3 = sideways). The sequence of flower orientations in the unstable treatment was assigned at the individual level, and the distribution of orientations across trials was approximately balanced across bees (Table S1).

Each bee performed three consecutive foraging trials (T1–T3) within a single day. A foraging trial was defined as a trip from the colony to the flight cage and back. Bees were considered to have stopped foraging if they returned to the corridor at the entrance of the flight cage to return to the colony by themselves or if they did not land on or approach a flower for more than 5 minutes (Russell et al., 2016).

Of 101 bees pre-trained to collect pollen, 37 (36.6%) returned and completed the baseline preference test (P1). Five of these did not return for subsequent trials and were excluded from further analyses. The analyses of baseline preference (P1) and experimental trials (T1–T3) include 32 bees (16 stable, 16 unstable), yielding 96 bee–trial observations. Post-trial preference (P2) was obtained for 31 of these individuals.

### Pollen load collection and measurement

At the end of each foraging trial, returning bees were intercepted in the corridor between the experimental flight cage and the foraging arena. A pollen load was removed from one randomly selected hind leg (corbicula) using fine forceps. Only one pollen load was removed to avoid disrupting subsequent foraging behaviour (Raine & Chittka, 2007a).

Pollen loads were weighed to the nearest milligram using an analytical balance. Because artificial flowers did not provide nectar, pollen load mass was used as a proxy for the amount of pollen collected (Raine & Chittka, 2007a).

Pollen load could not be measured in a subset of foraging trials (n = 37 across all trials and preference tests; 12 stable, 25 unstable), i.e. when bees returned without a recoverable pollen load, indicating negligible or no pollen collection during the trial. These observations were retained in the dataset and assigned a pollen load weight of zero for analysis.

### Behavioural data extraction

Flower visitation sequences were recorded live during experiments and compiled as digitised behavioural records for each bee and trial. From these records, we calculated the total number of flower visits, the number of unique flowers visited (floral coverage), the number of revisits to the same flower, revisit proportion (revisits / total visits), and unique-flower proportion (unique flowers / total visits). Trial duration was extracted from video recordings using BORIS (Friard & Gamba, 2016) and was defined as the time elapsed between the first and last flower visit within a trial. Sonication behaviour was also scored live during experiments as a binary variable (present/absent) based on auditory cues generated during visits to artificial flowers.

### Morphological measurements

To account for potential effects of body size, intertegular span (ITS) and body mass were measured for each bee after completion of the experiment.

### Ethical Note

This study involved commercially obtained *Bombus terrestris*, an invertebrate species not covered by the UK Animals (Scientific Procedures) Act 1986. Because the study did not involve protected animals under this legislation, formal approval from an institutional animal ethics committee was not required. Colonies were maintained in their original nest boxes in a climate-controlled laboratory maintained at 25 °C (relative humidity approximately 45–55%), with *ad libitum* access to sugar solution and pollen outside experimental foraging trials. Experimental procedures were designed to minimise disturbance to the bees. Individuals foraged voluntarily throughout the experiments and were handled only for individual identification and the removal of a single pollen load from one randomly selected corbicula at the end of each foraging trial, thereby minimising disruption to subsequent foraging behaviour. Bees were returned to their colonies after each trial. Following completion of the experiments, colonies were euthanised by freezing at −20 °C. The study was conducted in accordance with the ASAB/ABS Guidelines for the ethical treatment of nonhuman animals in behavioural research and teaching (ASAB Ethical Committee/ABS Animal Care Committee, 2025).

### Data analysis

All analyses were conducted in R (v4.4.2; (R Core Team, 2024). Mixed-effects models were fitted using the *lme4* package (Bates et al., 2015), including linear and generalized linear formulations where appropriate, and negative binomial models were fitted using *glmmTMB* (Brooks et al., 2017) for overdispersed count data. Model assumptions were assessed using simulation-based residual diagnostics implemented in *DHARMa* (Hartig, 2024).

Repeated measurements across trials were modelled using bee identity as a random intercept. Colony was included as a fixed effect, rather than a random effect, because only three colonies were sampled.

Baseline flower orientation preference (P1), prior to treatment exposure, was analysed separately using a negative binomial generalized linear mixed-effects model testing for differences in visitation among flower orientations, with bee identity included as a random intercept. Analyses were restricted to individuals retained in the main experiment.

To assess behavioural responses to environmental instability during the treatment phase (T1–T3), mixed-effects models were used to analyse revisitation behaviour, floral coverage (number of unique flowers visited), sonication behaviour, pollen collection success, pollen load, and trial duration. Fixed effects included treatment, trial, colony, and their interactions where biologically relevant. Behavioural predictors including floral coverage, visitation effort (total flower visits), revisitation probability, and sonication behaviour were subsequently incorporated into additional models to examine how different components of behavioural organisation related to pollen acquisition and time investment.

Additional analyses examined foraging efficiency (FE), calculated as estimated total pollen load collected divided by trial duration. Because unsuccessful pollen collection generated structural zeros, FE analyses were restricted to observations with non-zero pollen loads. During T1–T3, mixed-effects models included treatment, trial, their interaction, and colony as fixed effects and bee identity as a random intercept. For the final preference test (P2), FE was analysed using linear models including treatment and colony as fixed effects.

To assess persistent carry-over effects, data from the final preference test (P2) were analysed using linear or generalized linear models including treatment and colony as fixed effects. Responses included pollen load, pollen collection success, revisitation behaviour, floral coverage, visitation effort, trial duration, and sonication behaviour. Additional analyses restricted to successful pollen collectors were conducted to distinguish treatment effects on pollen collection success from effects on pollen load, bout duration, and foraging efficiency conditional on successful collection.

For mixed-effects and generalized linear mixed-effects models, significance of fixed effects was assessed using Type III Wald chi-square tests implemented in the *car* package (Fox & Weisberg, 2019). For linear models, significance was assessed using Type III F-tests.

Additional analyses examining relationships among revisitation behaviour and floral coverage, exploratory analyses of pollen extraction, orientation-specific carry-over during the post-trial preference test (P2) and morphology-augmented versions of the main models are described in Supplementary Methods S1–S3 and Table S2.

## Results

### 1. Baseline orientation preference (P1)

There was no strong evidence that flower visitation frequency differed across flower orientations during the initial preference test (n=32 bees) (Fig. S1), although there was a weak trend towards upward-facing flowers (χ²(2) = 5.66, *p* = 0.059). A non-parametric Friedman test on within-individual visit proportions likewise showed no significant effect of flower orientation on visitation in the initial test (χ²(2) = 2.09, *p* = 0.352). Together, these results indicate no clear baseline preference for flower orientation prior to experimental manipulation.

### 2. Environmental instability reorganised foraging behaviour

Environmental instability altered flower revisitation behaviour across successive foraging trials (Fig. 2A). Model-estimated revisit proportions were similar between treatments in the first trial (stable: 0.44; unstable: 0.42) but diverged across subsequent trials. Under stable conditions, revisit proportions remained relatively constant across trials (T1: 0.44, T2: 0.44, T3: 0.48), whereas under unstable conditions they declined progressively (T1: 0.42, T2: 0.35, T3: 0.22). Consistent with this pattern, the treatment × trial interaction was marginal (χ²(2) = 5.41, p = 0.067), while there was a significant main effect of treatment (χ²(1) = 4.58, *p* = 0.032), but no strong main effect of trial (χ²(1) = 2.57, *p* = 0.276).

**Figure 2.**
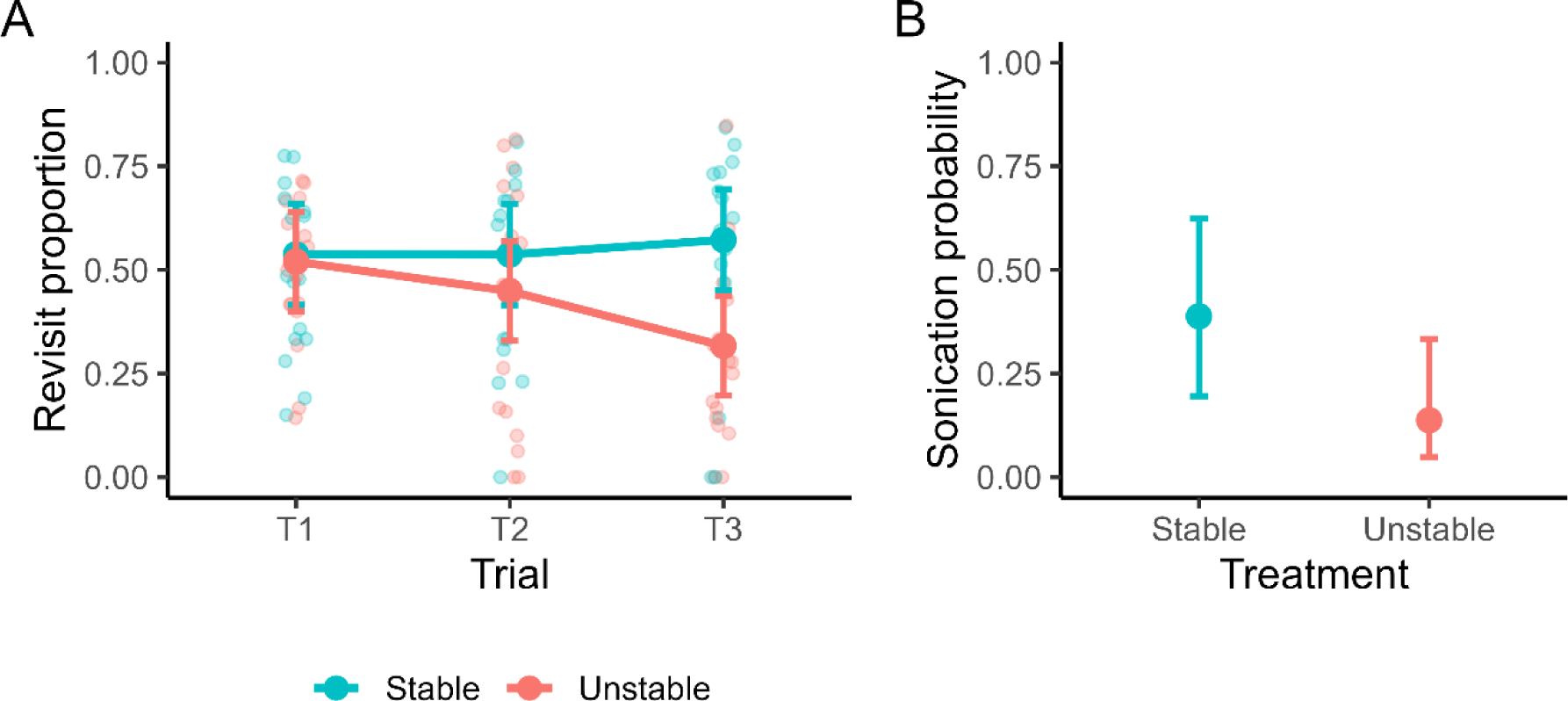
Environmental instability altered revisitation and sonication behaviour during pollen foraging. (A) Revisit proportion across three successive foraging trials under stable and unstable floral orientation treatments. Points and error bars show model-estimated means ± 95% confidence intervals from the factor-coded mixed-effects model; faint points show individual observations. Revisitation probability remained relatively constant under stable conditions but declined across successive trials under unstable conditions. (B) Model-estimated probability of sonication under stable and unstable floral orientation treatments. Points and error bars show estimated marginal means ± 95% confidence intervals from the fitted mixed-effects model. Bees exposed to unstable floral orientations exhibited lower probabilities of sonication than bees foraging under stable conditions.

Environmental instability also altered flower-handling behaviour (Fig. 2B). Sonication probability differed significantly between treatments (χ²(1) = 7.51, *p* = 0.006), with bees foraging under stable conditions exhibiting substantially higher probabilities of sonication than bees exposed to unstable floral conditions (stable: 0.388 [95% CI: 0.195–0.623]; unstable: 0.137 [95% CI: 0.048– 0.333]). This effect remained significant when accounting for variation among foraging bouts (χ²(1) = 5.99, *p* = 0.014). In contrast, effects of trial were weak and inconsistent across models (e.g., χ²(1) = 5.55, *p* = 0.018 in the primary model, but χ²(1) = 1.54, *p* = 0.215 when accounting for bout-level variation). Sonication behaviour additionally showed strong temporal structure within foraging bouts, being strongly predicted by behaviour on the previous visit (χ²(1) = 270.68, *p* < 0.001) and varying non-linearly across visit sequences (linear: χ²(1) = 117.34, *p* < 0.001; quadratic: χ²(1) = 73.96, *p* < 0.001; Fig. S3).

Together, these results demonstrate that environmental instability altered both how bees distributed visits among flowers and how they interacted with flowers during pollen collection.

### 3. Different components of behavioural organisation predicted pollen acquisition and time investment

To determine whether different measures of foraging behaviour captured similar or distinct aspects of behavioural organisation, we examined the relationships among revisitation behaviour, floral coverage, and visitation effort. Floral coverage, defined as the number of unique flowers visited within a foraging trial, represents the extent to which individuals explored available floral resources. Revisitation probability and floral coverage were moderately correlated across individuals and trials (see Supplementary Methods S1; Pearson’s r = 0.60, 95% CI [0.45, 0.71], p < 0.001). However, this association was driven primarily by variation in total visit number rather than a direct relationship between revisitation and floral coverage.

Revisitation probability was strongly associated with total visit number (χ²(1) = 295.74, p < 0.001; Fig. 3A), indicating that higher revisit rates primarily reflected increased visitation effort. In contrast, when revisitation probability and total visit number were included in the same model, floral coverage was significantly predicted by total visit number (χ²(1) = 8.89, p = 0.003), but not by revisitation probability (χ²(1) = 1.10, p = 0.294; Fig. 3B). There was also no evidence that floral coverage differed between treatments or across trials (all p > 0.70; Fig. S2). Together, these results demonstrate that revisitation behaviour, floral coverage, and visitation effort capture distinct components of behavioural organisation rather than interchangeable measures of exploration.

**Figure 3.**
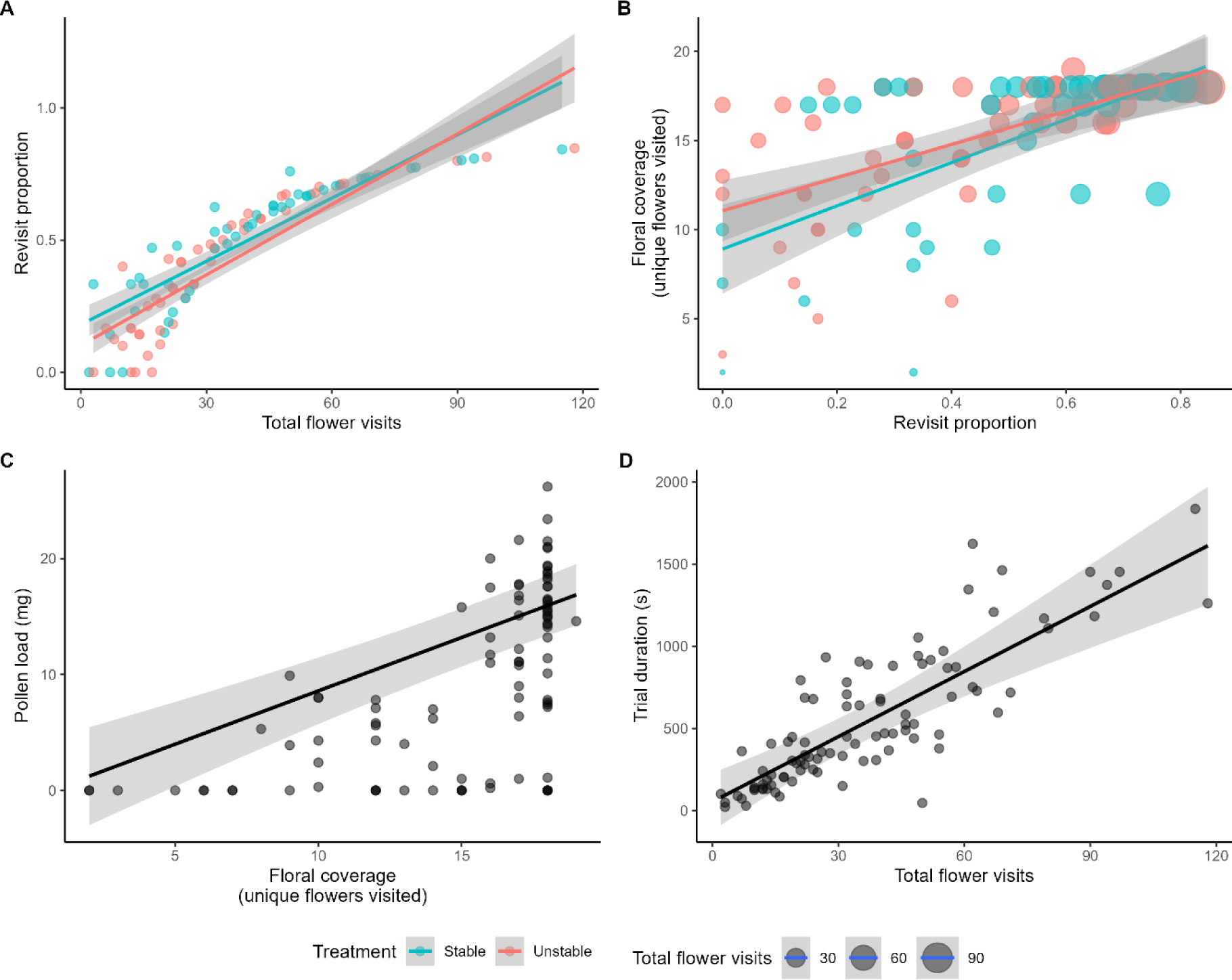
Different components of behavioural organisation predicted pollen acquisition and time investment. (A) Relationship between revisit proportion and total flower visits, showing a strong positive association consistent with revisitation primarily reflecting visitation effort. Coloured lines indicate fitted relationships for bees foraging under stable and unstable conditions. (B) Relationship between revisit proportion and floral coverage (number of unique flowers visited), illustrating that revisitation and floral coverage capture related but distinct components of foraging behaviour. Point size is proportional to total flower visits. (C) Relationship between floral coverage and pollen load, showing that bees visiting more unique flowers collected larger pollen loads. (D) Relationship between total flower visits and trial duration, showing that time investment increased with visitation effort. Points represent individual observations, and lines show model fits ± 95% confidence intervals.

We next asked whether these different behavioural components predicted different foraging outcomes. Among bees that collected pollen, pollen acquisition was primarily driven by floral coverage rather than revisitation behaviour or visitation effort. Bees that visited more unique flowers collected significantly larger pollen loads overall (χ²(1) = 12.05, p < 0.001; Fig. 3C). In contrast, neither total visit number (χ²(1) = 0.97, p = 0.324) nor revisitation probability (χ²(1) = 1.86, p = 0.173) significantly predicted pollen load once floral coverage was accounted for.

Time investment showed a different pattern. Trial duration was primarily driven by visitation effort, with the number of visits having a strong positive effect on trial duration (χ²(1) = 41.73, p < 0.001; Fig. 3D). In contrast, neither floral coverage (χ²(1) = 1.36, p = 0.244) nor revisitation probability (χ²(1) = 0.01, p = 0.917) significantly influenced trial duration. There was also no evidence for effects of trial (χ²(2) = 0.06, p = 0.970) or treatment (χ²(1) = 0.24, p = 0.627), while colony identity contributed significantly to variation in trial duration (χ²(2) = 9.69, p = 0.008).

Together, these results demonstrate that different components of behavioural organisation predicted different foraging outcomes. Floral coverage was associated with pollen acquisition, whereas visitation effort primarily predicted the time spent foraging.

### 4. Environmental instability had limited immediate effects on pollen acquisition

During T1-T3, treatment did not affect the probability that bees returned with any recoverable pollen (χ²(1) = 0.62, p = 0.432), and pollen collection success did not vary significantly across trials (χ²(2) = 4.57, p = 0.102). However, bees exposed to unstable floral conditions tended to collect smaller pollen loads than bees foraging under stable conditions, although this effect was only marginally supported statistically (χ²(1) = 3.73, p = 0.054). Pollen load did not vary across trials (χ²(2) = 0.21, p = 0.902), while colony identity contributed significantly to variation in pollen collection (χ²(2) = 14.25, p < 0.001). Sonication behaviour was not a significant predictor of pollen load when included in the model (χ²(1) = 1.17, *p* = 0.280), indicating that differences in sonication behaviour did not explain treatment differences in pollen load. Among bees that successfully collected pollen, foraging efficiency varied across treatments and trials (Treatment × Trial: χ²(2) = 8.80, p = 0.012). Post hoc comparisons revealed no treatment differences in T1 (p = 0.222) or T2 (p = 0.733), but in T3, bees that successfully collected pollen in the unstable treatment exhibited higher foraging efficiency than those in the stable treatment (p = 0.029). Within the unstable treatment group, foraging efficiency was also higher in T3 than in T1 (p = 0.034). Examination of the underlying components of foraging efficiency indicated that this difference was associated primarily with unstable bees having shorter foraging bouts (F_1,18_ = 5.68, p = 0.028), while pollen load mass did not differ significantly between treatments (F_1,19_ = 3.14, p = 0.092).

Thus, despite clear effects of instability on foraging behaviour and organisation, its immediate effects on pollen acquisition were comparatively weak. Although treatment did not affect the probability of successful pollen collection, successful bees in the unstable treatment exhibited higher foraging efficiency by the final treatment trial, driven primarily by reduced foraging time rather than increased pollen acquisition.

### 5. Environmental instability generated carry-over effects on pollen acquisition

Prior exposure to environmental instability produced persistent effects on pollen acquisition that became apparent during the post-treatment preference test. In the final preference test (P2), bees from both treatments were presented with flowers of all three orientations simultaneously. Each flower contained sufficient pollen (20 mg/flower) that bees could theoretically collect a full pollen load by visiting flowers of a single orientation alone. In this scenario, bees previously exposed to unstable conditions collected significantly less pollen overall than bees previously exposed to stable conditions (F_1,27_ = 7.68, p = 0.009; Fig. 4A) and were also less likely to return with any recoverable pollen load than bees from the stable treatment (χ²(1) = 10.53, p = 0.001; Fig. 4B), indicating that differences between treatment groups persisted during the common post-treatment preference test.

**Figure 4.**
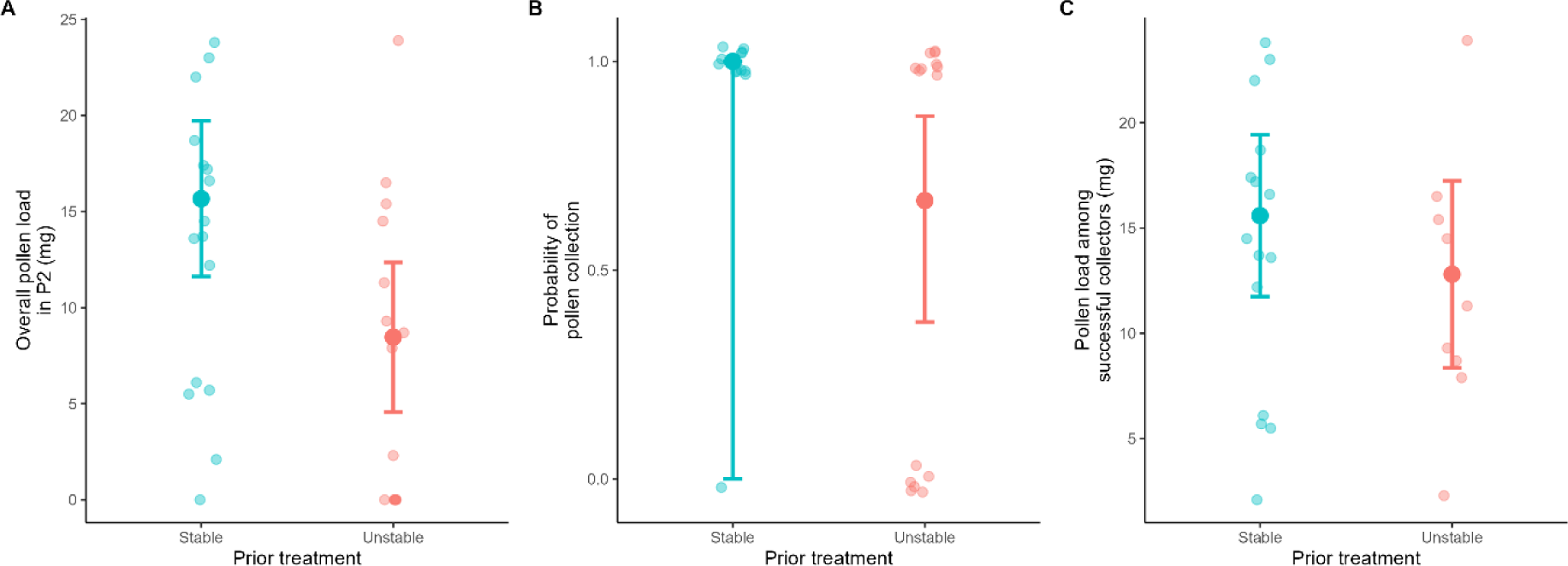
Carry-over effects of environmental instability on pollen acquisition. (A) Overall pollen load collected during the final preference test (P2), in which bees from both treatments were presented with the same array containing flowers of all three orientations and abundant pollen rewards. Bees previously exposed to unstable floral conditions collected less pollen overall than bees previously exposed to stable conditions. (B) Probability of collecting any pollen during P2, showing that bees from the unstable treatment were less likely to return with recoverable pollen. (C) Pollen load among successful pollen collectors only, showing that pollen load did not differ between treatments once bees successfully collected pollen. Points represent individual observations, and large points with error bars show model-estimated means ± 95% confidence intervals. Together, these results indicate that the reduction in overall pollen acquisition following prior exposure to instability was primarily driven by a lower probability of successful pollen collection rather than reduced pollen loads among successful foragers.

To determine whether this reduction reflected lower pollen loads among successful foragers, we repeated the analysis using only bees that returned with non-zero pollen loads. Among successful pollen collectors, pollen load did not differ between treatments (F_1,21_ = 1.06, p = 0.316; Fig. 4C). Additional analyses likewise revealed no treatment differences in bout duration (F_1,21_ = 0.94, p = 0.344) or foraging efficiency (F_1,21_ = 1.65, p = 0.213) among successful pollen collectors.

Thus, the overall reduction in pollen acquisition following exposure to instability was driven primarily by a lower probability of successful pollen collection rather than by reduced pollen loads, reduced foraging efficiency, or longer foraging bouts among successful foragers.

We next asked whether the reduced pollen acquisition observed in bees previously exposed to unstable conditions could be explained by variation in behavioural organisation. Bees previously exposed to stable conditions exhibited higher revisitation rates than bees from the unstable treatment (F_1,27_ = 12.77, p = 0.001), whereas treatment effects on total visit number were weaker (F_1,27_ = 3.33, p = 0.079) and floral coverage did not differ significantly between treatments (F_1,27_ = 2.28, p = 0.143). However, neither revisitation behaviour, visitation effort, nor floral coverage significantly predicted pollen load in P2 (all p > 0.35). Moreover, treatment remained a significant predictor of pollen load after accounting for floral coverage (F_1,26_ = 6.27, p = 0.019) and visitation effort (F_1,26_ = 5.57, p = 0.026). Treatment also remained a significant predictor of pollen collection success after accounting for floral coverage and revisitation behaviour (χ²(1) = 4.44, p = 0.035).

We additionally tested whether persistent treatment differences in pollen acquisition were associated with variation in sonication behaviour. There was no evidence that sonication differed between treatments in P2 (F_1,27_ = 1.29, p = 0.266), and sonication behaviour did not predict pollen load when included alongside treatment and colony (F_1,26_ = 1.63, p = 0.213). In contrast, treatment remained a significant predictor of pollen load (F_1,26_ = 9.38, p = 0.005).

Exploratory analyses comparing bees that did and did not return with recoverable pollen revealed no differences in visitation effort, floral coverage, sonication behaviour or foraging duration after accounting for treatment (all p > 0.16).

Finally, treatment differences in pollen collection during P2 were not accompanied by orientation-specific visitation biases. Bees did not preferentially visit either the orientation experienced throughout training (stable treatment) or the final orientation experienced before P2 (unstable treatment; Fig. S4).

Together, these results demonstrate that prior exposure to environmental instability had persistent effects on pollen acquisition during the common post-treatment preference test. These carry-over effects were expressed primarily through a reduced probability of successful pollen collection and were not explained by differences in foraging efficiency, visitation effort, floral coverage, revisitation behaviour, sonication behaviour or flower orientation visitation biases.

## Discussion

Environmental predictability is expected to shape how animals organise behaviour because it determines the value of information acquired through previous experience (Dall et al., 2005; McNamara & Dall, 2010; Stephens, 1989). Here, we experimentally manipulated environmental predictability through repeated changes in the floral orientation encountered by pollen-foraging bumblebees between foraging trips. We found that environmental instability reorganised foraging behaviour and generated persistent reductions in subsequent pollen acquisition. Strikingly, these costs emerged despite relatively weak immediate effects on pollen collection, suggesting that behavioural flexibility buffered short-term performance while masking delayed consequences of environmental instability.

The most important finding of this study was the emergence of carry-over effects during the post-treatment preference test. Because both treatment groups experienced identical foraging conditions during this final test, it provided a common assay of whether prior exposure to environmental instability had persistent effects after the environmental differences themselves between the groups had been removed. During this final test, both treatment groups were presented with the same array containing flowers of all three orientations and abundant pollen rewards, yet bees previously exposed to unstable floral orientations between foraging trials collected less pollen overall and were substantially less likely to return with recoverable pollen. Importantly, this treatment effect disappeared when analyses were restricted to successful pollen collectors who returned with quantifiable loads, indicating that instability primarily influenced the probability of successful pollen acquisition rather than the amount of pollen collected once acquisition occurred. Because the behavioural variables measured here did not account for this treatment effect, the mechanism underlying the reduced probability of successful pollen acquisition remains unresolved. One possibility is that repeated exposure to environmental instability altered motivational state, reducing the persistence or investment with which bees harvested pollen. Alternatively, instability may have affected finer-scale aspects of pollen extraction or handling that were not captured by our behavioural observations.

Previous environmental conditions are increasingly recognised as important determinants of subsequent behaviour and performance across a wide range of ecological contexts (Harrison et al., 2011; O’connor et al., 2014). Experimental studies have further demonstrated that prior exposure to environmental variability can alter subsequent memory use, exploratory behaviour and information processing even after the original environmental conditions are removed (Abts & Dunlap, 2022; Dunlap & Stephens, 2012; Young & Dyer, 2022). Consistent with this broader literature, our results demonstrate that exposure to environmental instability can generate carry-over effects on pollinator foraging behaviour that persist after environmental differences themselves have disappeared.

The delayed nature of these effects is particularly noteworthy because bees appeared remarkably resilient during the period of instability itself. Environmental instability altered both revisitation behaviour and flower-handling behaviour, yet immediate effects on pollen acquisition were comparatively weak. Bees exposed to unstable conditions were no less likely to collect pollen and showed only a marginal tendency to return with smaller pollen loads. Moreover, by the final foraging trial, among successful pollen collectors, bees exposed to instability collected similar pollen loads to those in the stable treatment but completed foraging bouts more quickly, resulting in higher foraging efficiency. Together, these findings suggest that bees adjusted their behaviour in ways that largely maintained short-term performance despite repeated environmental change. Bumblebees are well known for their capacity to modify behaviour in response to changing environmental conditions, including reversals in reward contingencies, changing cue reliability and variation in resource quality (Chittka, 1998; Dunlap et al., 2017; Raine & Chittka, 2012; Strang & Sherry, 2014). More generally, behavioural flexibility is widely considered an important mechanism through which organisms cope with environmental variability (Dall et al., 2005; Leimar et al., 2024; McNamara & Dall, 2010). Our results support this view but also suggest an important qualification. Behavioural adjustments appeared sufficient to maintain short-term performance under unstable conditions, yet delayed costs emerged later. Behavioural flexibility therefore did not eliminate the consequences of instability; instead, it appeared to postpone their expression.

This distinction may have broader implications for understanding biological responses to environmental change. Many studies assess the consequences of environmental variability using immediate behavioural or performance responses. However, experimental work in both vertebrates and insects suggests that previous environmental conditions can influence subsequent behaviour long after the original experience has ended (Abts & Dunlap, 2022; Dunlap & Stephens, 2012; Young & Dyer, 2022). Our findings extend this perspective by demonstrating that behavioural resilience during periods of instability does not necessarily imply an absence of costs. If organisms compensate behaviourally while environmental conditions are changing, assessments based solely on concurrent measures may underestimate the biological consequences of environmental variability.

One of the clearest behavioural responses to instability was a progressive reduction in revisitation behaviour across successive foraging bouts. Bees exposed to stable conditions maintained relatively constant revisit rates, whereas individuals exposed to changing floral orientations revisited flowers less frequently over time. Repeated revisitation is a common feature of route-based foraging and trapline formation, in which pollinators repeatedly exploit familiar resource locations using learned information (Lihoreau et al., 2012; Saleh & Chittka, 2007; Thomson, 1996). Although our experimental design was not intended to test trapline formation directly, the observed decline in within-bout revisitation suggests that repeated environmental instability altered how bees organised their foraging routes. One possibility is that repeated changes in floral orientation reduced the reliability of information acquired during previous foraging bouts, diminishing the benefits of repeatedly exploiting familiar flowers. This interpretation is consistent with theoretical expectations that animals should adjust their reliance on previously acquired information according to the predictability of environmental states and the expected value of that information for future decisions (Dall et al., 2005; McNamara & Dall, 2010; Stephens, 1989).

Importantly, reduced revisitation did not reflect reduced exploration of available resources. Floral coverage remained remarkably stable across treatments, indicating that bees continued to sample a similar proportion of available flowers despite changes in revisit behaviour. Revisitation behaviour, visitation effort and floral coverage therefore captured distinct aspects of foraging organisation rather than alternative measures of a single exploratory axis. This distinction proved important because these behavioural variables predicted different components of foraging performance. Floral coverage was positively associated with pollen acquisition, whereas visitation effort primarily influenced bout duration. Environmental instability therefore altered how bees organised foraging behaviour rather than simply increasing or decreasing overall foraging activity. Similar dissociations between movement patterns and resource exploitation have been documented in studies of spatial learning and route optimisation, where animals modify movement structure while maintaining comparable levels of resource sampling (Lihoreau et al., 2010; Woodgate et al., 2017).

Environmental instability also reduced the probability of sonication. Sonication is a specialised pollen-foraging behaviour associated with pollen extraction from flowers possessing poricidal anthers (Buchmann, 1983; De Luca & Vallejo-Marín, 2013), and its expression can be influenced by both floral traits and previous experience (Morgan et al., 2016). The reduction in sonication observed under unstable conditions therefore suggests that instability influenced not only movement decisions but also flower-handling behaviour. However, sonication did not predict pollen acquisition and therefore cannot explain the treatment differences observed here. Rather, instability appears to have affected multiple components of behavioural organisation simultaneously, only some of which translated directly into measurable differences in resource collection.

The mechanisms underlying the observed carry-over effects remain unresolved. In the post-treatment preference test (P2), persistent treatment differences could not be explained by revisitation behaviour, visitation effort, floral coverage, sonication behaviour, body size or flower orientation visitation biases. Exploratory analyses further indicated that, although pollen removal from flowers was closely associated with pollen collection during the treatment phase, this relationship disappeared during the final preference test. Thus, the persistent reduction in pollen acquisition following exposure to instability was unlikely to arise solely from differences in pollen extraction and instead appears to have emerged during subsequent stages of pollen collection, such as the efficiency of pollen handling, grooming or transfer of pollen into the corbiculae, suggesting that environmental instability influenced finer-scale aspects of pollen foraging that were not captured by the behavioural variables measured here.

One possibility is that repeated environmental instability altered how bees used information acquired through previous experience. Information-use theory predicts that learning should be most beneficial when environmental conditions are sufficiently predictable that acquired information retains future value (Dall et al., 2005; McNamara & Dall, 2010; Stephens, 1989). Similarly, adaptive memory theory predicts that retention should depend on the temporal structure of environmental change, because information becomes less useful when environmental states change rapidly (Anderson & Schooler, 1991; Dunlap & Stephens, 2012). By repeatedly altering floral orientation while maintaining reward availability, our manipulation specifically reduced the predictability of a cue associated with resource exploitation without altering the rewards themselves. Under such conditions, individuals may be required to repeatedly update behavioural responses, potentially generating costs that are not immediately expressed in resource acquisition. Although our experiment did not directly measure learning, memory or cognition, the persistence of treatment effects after behavioural differences largely disappeared is consistent with the possibility that instability altered how previously acquired information was subsequently used during foraging.

Our findings may also contribute to a growing effort to understand how behavioural processes mediate responses to environmental variability. Environmental change is increasingly altering the spatial, temporal and informational structure of ecological systems through habitat modification, altered species interactions and climate-driven shifts in resource availability (Burkle et al., 2013; Memmott et al., 2007; Miller-Struttmann et al., 2015). Considerable attention has therefore focused on whether behavioural flexibility allows organisms to cope with changing conditions. The present results suggest a more nuanced perspective. Behavioural flexibility may enable organisms to maintain performance while conditions are changing, yet delayed costs may still emerge if environmental instability reduces the future value of information acquired through experience. Understanding the consequences of environmental change may therefore require considering not only immediate behavioural responses, but also how changing environments influence the long-term usefulness of learned information.

Our manipulation focused on a single floral trait under controlled laboratory conditions, and future work is needed to determine whether similar effects arise under more complex ecological conditions and in response to other forms of environmental instability. Environmental predictability can be manipulated through multiple dimensions, including reward variance, state persistence and cue reliability (Dall et al., 2005; Koops, 2004; Stephens, 1989). Our experiment specifically altered the predictability of a floral cue while maintaining reward availability, however future studies should investigate whether different forms of environmental uncertainty produce similar behavioural and carry-over effects. Future studies could also investigate whether behavioural responses to environmental instability influence plant reproductive success by altering pollen transfer dynamics.

In conclusion, environmental instability altered the organisation of pollen-foraging behaviour and generated persistent reductions in pollen collection success that were evident during the common post-treatment preference test. These delayed effects were not explained by differences in visitation effort, resource coverage, sonication behaviour, morphology, pollen extraction or flower orientation visitation biases, suggesting that instability influenced aspects of behavioural performance that extend beyond commonly measured foraging metrics. More broadly, our results demonstrate that behavioural flexibility can maintain short-term performance under changing conditions while masking delayed costs that emerge later. Understanding how organisms respond to increasingly variable environments may therefore require considering not only how behaviour changes during environmental instability, but also the longer-term consequences of relying on information whose value is continually changing.

## Data Availability

The data and R code supporting the findings of this study are provided as Supplementary Material.

## Acknowledgements

We thank Becca Gorham for assisting with data collection, as well as Lucy Unwin, Scarlett Dowling, and Angela Wangtapan for assisting with video coding and data digitisation. We also thank Avery Russell for advice with pollen supply and bumblebee supplier. This study was supported by a UKRI Future Leaders Fellowship (MR/T021691/1).

## Notes

### Competing Interest Statement

The authors have declared no competing interest.

## References

Abts, B. J., & Dunlap, A. S. (2022). Memory and the value of social information in foraging bumble bees. Learning and Behavior, 50(3), 317–328. 10.3758/S13420-022-00528-2/FIGURES/5

Anderson, J. R., & Schooler, L. J. (1991). Reflections of the Environment in Memory. Psychological Science, 2(6), 396–408. 10.1111/J.1467-9280.1991.TB00174.X/ASSET/8BFCA03F-B12C-401C-8F32-8FBE280DD366/ASSETS/IMAGES/LARGE/10.1111_J.1467-9280.1991.TB00174.X-FIG14.JPG

Armbruster, W. S., & Muchhala, N. (2020). Floral reorientation: the restoration of pollination accuracy after accidents. New Phytologist, 227(1), 232–243. 10.1111/nph.16482

ASAB Ethical Committee/ABS Animal Care Committee. (2025). Guidelines for the ethical treatment of nonhuman animals in behavioural research and teaching. Animal Behaviour, 219, 123065. 10.1016/S0003-3472(24)00376-2

Bates, D., Mächler, M., Bolker, B. M., & Walker, S. C. (2015). Fitting linear mixed-effects models using lme4. Journal of Statistical Software, 67(1), 1–48. 10.18637/JSS.V067.I01

Brooks, M., Kristensen, K., Benthem, K. van, Magnusson, A., Berg, C., Nielsen, A., Skaug, H., Mächler, M., & Bolker, B. (2017). glmmTMB balances speed and flexibility among packages for zero-inflated generalized linear mixed modeling. The R Journal.

Buchmann, S. L. (1983). Buzz pollination in angiosperms. In C. E. Jones & R. J. Little (Eds.), Handbook of experimental pollination biology (pp. 73–113). Van Nostrand Reinhold Company.

Burkle, L. A., Marlin, J. C., & Knight, T. M. (2013). Plant-pollinator interactions over 120 years: Loss of species, co-occurrence, and function. Science, 340(6127), 1611–1615. 10.1126/SCIENCE.1232728/SUPPL_FILE/BURKLE.SM.PDF

Chittka, L. (1998). Sensorimotor learning in bumblebees: long-term retention and reversal training. Journal of Experimental Biology, 201(4), 515–524. 10.1242/JEB.201.4.515

Chittka, L., & Thomson, J. D. (2001). Cognitive ecology of pollination: animal behaviour and floral evolution (L. Chittka & J. D. Thomson, Eds.). Cambridge University Press. https://books.google.co.uk/books?id=g2Km4B6n-mQC&dq=Chittka+L,+Thomson+JD+(Eds.),+Cognitive+Ecology+of+Pollination+-+Animal+Behavior+and+Floral+Evolution,+Cambridge+University+Press,+Cambridge,++pp.+191-213.+&lr=&hl=fr&source=gbs_navlinks_s

Chittka, L., Thomson, J. D., & Waser, N. M. (1999). Flower constancy, insect psychology, and plant evolution. Naturwissenschaften 1999 86:8, 86(8), 361–377. 10.1007/S001140050636

Dall, S., Giraldeau, L., Olsson, O., McNamara, J., & Stephens, D. (2005). Information and its use by animals in evolutionary ecology. Trends in Ecology & Evolution, 20(4), 187–193. 10.1016/j.tree.2005.01.010

De Luca, P. A., & Vallejo-Marín, M. (2013). What’s the ‘buzz’ about? The ecology and evolutionary significance of buzz-pollination. Current Opinion in Plant Biology, 16(4), 429–435. 10.1016/J.PBI.2013.05.002

Dunlap, A. S., Papaj, D. R., & Dornhaus, A. (2017). Sampling and tracking a changing environment: Persistence and reward in the foraging decisions of bumblebees. Interface Focus, 7(3). 10.1098/RSFS.2016.0149/64158

Dunlap, A. S., & Stephens, D. W. (2012). Tracking a changing environment: optimal sampling, adaptive memory and overnight effects. Behavioural Processes, 89(2), 86–94. 10.1016/J.BEPROC.2011.10.005

Fox, J., & Weisberg, S. (2019). An R Companion to Applied Regression (Third). Sage.

Friard, O., & Gamba, M. (2016). BORIS: a free, versatile open-source event-logging software for video/audio coding and live observations. Methods in Ecology and Evolution, 7(11), 1325–1330. 10.1111/2041-210X.12584

Grüter, C. (2026). Flower constancy in pollinators: a bouquet of agendas shapes interactions among mutualistic partners. Proceedings of the Royal Society B: Biological Sciences, 293(2064), 20252103. 10.1098/rspb.2025.2103

Harrison, X. A., Blount, J. D., Inger, R., Norris, D. R., & Bearhop, S. (2011). Carry-over effects as drivers of fitness differences in animals. Journal of Animal Ecology, 80(1), 4–18. 10.1111/J.1365-2656.2010.01740.X

Hartig, F. (2024). DHARMa: residual diagnostics for hierarchical (multi-level / mixed) regression models.

Jirgal, N., & Ohashi, K. (2023). Effects of floral symmetry and orientation on the consistency of pollinator entry angle. The Science of Nature, 110(3), 19. 10.1007/s00114-023-01845-w

Julesz, B. (1962). Visual pattern discrimination. IEEE Transactions on Information Theory, 8(2), 84–92. 10.1109/TIT.1962.1057698

Koops, M. A. (2004). Reliability and the value of information. Animal Behaviour, 67(1), 103–111. 10.1016/j.anbehav.2003.02.008

Leimar, O., Quiñones, A. E., & Bshary, R. (2024). Flexible learning in complex worlds. Behavioral Ecology, 35(1), 1–12. 10.1093/BEHECO/ARAD109

Lihoreau, M., Chittka, L., & Raine, N. E. (2010). Travel optimization by foraging bumblebees through readjustments of traplines after discovery of new feeding locations. The American Naturalist, 176(6), 744–757. 10.1086/657042

Lihoreau, M., Raine, N. E., Reynolds, A. M., Stelzer, R. J., Lim, K. S., Smith, A. D., Osborne, J. L., & Chittka, L. (2012). Radar tracking and motion-sensitive cameras on flowers reveal the development of pollinator multi-destination routes over large spatial scales. PLoS Biology, 10(9), e1001392. 10.1371/journal.pbio.1001392

Makino, T. T., & Thomson, J. D. (2012). Innate or learned preference for upward-facing flowers?: implications for the costs of pendent flowers from experiments on captive bumble bees. Journal of Pollination Ecology, 9, 79–84. 10.26786/1920-7603(2012)11

McNamara, J. M., & Dall, S. R. X. (2010). Information is a fitness enhancing resource. Oikos, 119(2), 231–236. 10.1111/J.1600-0706.2009.17509.X

McNamara, J. M., & Houston, A. I. (1987). Memory and the efficient use of information. Journal of Theoretical Biology, 125(4), 385–395. 10.1016/S0022-5193(87)80209-6

Memmott, J., Craze, P. G., Waser, N. M., & Price, M. V. (2007). Global warming and the disruption of plant– pollinator interactions. Ecology Letters, 10(8), 710–717. 10.1111/J.1461-0248.2007.01061.X

Miller-Struttmann, N. E., Geib, J. C., Franklin, J. D., Kevan, P. G., Holdo, R. M., Ebert-May, D., Lynn, A. M., Kettenbach, J. A., Hedrick, E., & Galen, C. (2015). Functional mismatch in a bumble bee pollination mutualism under climate change. Science, 349(6255), 1541–1544. 10.1126/SCIENCE.AAB0868/SUPPL_FILE/MILLER-STRUTTMANN-SM.PDF

Morgan, T., Whitehorn, P., Lye, G. C., & Vallejo-Marín, M. (2016). Floral sonication is an innate behaviour in bumblebees that can be fine-tuned with experience in manipulating flowers. Journal of Insect Behavior, 29(2), 233–241. 10.1007/S10905-016-9553-5/TABLES/1

Nakata, T., Rin, I., Yaida, Y. A., & Ushimaru, A. (2022). Horizontal orientation facilitates pollen transfer and rain damage avoidance in actinomorphic flowers of *Platycodon grandiflorus*. Plant Biology, 24(5), 798–805. 10.1111/plb.13414

Nevard, L., & Vallejo-Marín, M. (2022). Floral orientation affects outcross-pollen deposition in buzz-pollinated flowers with bilateral symmetry. American Journal of Botany, 109(10), 1568–1578. 10.1002/ajb2.16078

Nicholls, E., & Hempel de Ibarra, N. (2017). Assessment of pollen rewards by foraging bees. Functional Ecology, 31(1), 76–87. 10.1111/1365-2435.12778/SUPPINFO

O’connor, C. M., Norris, D. R., Crossin, G. T., & Cooke, S. J. (2014). Biological carryover effects: linking common concepts and mechanisms in ecology and evolution. Ecosphere, 5(3), 1–11. 10.1890/ES13-00388.1

R Core Team. (2024). R: a language and environment for statistical computing.

Raine, N. E., & Chittka, L. (2007a). Pollen foraging: Learning a complex motor skill by bumblebees (*Bombus terrestris*). Naturwissenschaften, 94(6), 459–464. 10.1007/S00114-006-0184-0/TABLES/1

Raine, N. E., & Chittka, L. (2007b). The Adaptive Significance of Sensory Bias in a Foraging Context: Floral Colour Preferences in the Bumblebee *Bombus terrestris*. PLOS ONE, 2(6), e556. 10.1371/JOURNAL.PONE.0000556

Raine, N. E., & Chittka, L. (2012). No trade-off between learning speed and associative flexibility in bumblebees: a reversal learning test with multiple colonies. PLOS ONE, 7(9), e45096. 10.1371/JOURNAL.PONE.0045096

Raine, N. E., Ings, T. C., Dornhaus, A., Saleh, N., & Chittka, L. (2006). Adaptation, genetic drift, pleiotropy, and history in the evolution of bee foraging behavior. Advances in the Study of Behavior, 36, 305–354. 10.1016/S0065-3454(06)36007-X

Ruan, C. J., & da Silva, J. A. T. (2011). Adaptive significance of floral movement. Critical Reviews in Plant Sciences, 30(4), 293–328. 10.1080/07352689.2011.587715

Russell, A. L., Leonard, A. S., Gillette, H. D., & Papaj, D. R. (2016). Concealed floral rewards and the role of experience in floral sonication by bees. Animal Behaviour, 120, 83–91. 10.1016/J.ANBEHAV.2016.07.024

Russell, A. L., & Papaj, D. R. (2016). Artificial pollen dispensing flowers and feeders for bee behaviour experiments. Journal of Pollination Ecology, 18(3), 13–22. http://hdl.handle.net/10150/621206

Saleh, N., & Chittka, L. (2007). Traplining in bumblebees (*Bombus impatiens*): A foraging strategy’s ontogeny and the importance of spatial reference memory in short-range foraging. Oecologia, 151(4), 719–730. 10.1007/S00442-006-0607-9/FIGURES/5

Sponsler, D., Iverson, A., & Steffan-Dewenter, I. (2023). Pollinator competition and the structure of floral resources. Ecography, 2023(9), e06651. 10.1111/ECOG.06651

Stephens, D. W. (1987). On economically tracking a variable environment. Theoretical Population Biology,32(1), 15–25. 10.1016/0040-5809(87)90036-0

Stephens, D. W. (1989). Variance and the Value of Information. The American Naturalist, 134(1), 128–140. 10.1086/284969

Stephens, D. W. (1991). Change, regularity, and value in the evolution of animal learning. Behavioral Ecology, 2(1), 77–89. 10.1093/beheco/2.1.77

Strang, C. G., & Sherry, D. F. (2014). Serial reversal learning in bumblebees (*Bombus impatiens*). Animal Cognition, 17(3), 723–734. 10.1007/S10071-013-0704-1/FIGURES/6

Thomson, J. D. (1996). Trapline foraging by bumblebees: I. Persistence of flight-path geometry. Behavioral Ecology, 7(2), 158–164. 10.1093/BEHECO/7.2.158

Vallejo-Marín, M. (2022). How and why do bees buzz? Implications for buzz pollination. Journal of Experimental Botany, 73(4), 1080–1092. 10.1093/JXB/ERAB428

Van Der Kooi, C. J., Kevan, P. G., & Koski, M. H. (2019). The thermal ecology of flowers. Annals of Botany, 124(3), 343–353. 10.1093/AOB/MCZ073

Van Doorn, W. G., & Van Meeteren, U. (2003). Flower opening and closure: a review. Journal of Experimental Botany, 54(389), 1801–1812. 10.1093/JXB/ERG213

Waser, N. M. (1986). Flower constancy: definition, cause, and measurement. The American Naturalist, 127(5), 593–603. 10.1086/284507

Woodgate, J. L., Makinson, J. C., Lim, K. S., Reynolds, A. M., & Chittka, L. (2017). Continuous radar tracking illustrates the development of multi-destination routes of bumblebees. Scientific Reports, 7(1), 17323. 10.1038/s41598-017-17553-1

Woodrow, C., Jafferis, N., Kang, Y., & Vallejo-Marín, M. (2024). Buzz-pollinating bees deliver thoracic vibrations to flowers through periodic biting. Current Biology, 34(18), 4104–4113.e3. 10.1016/j.cub.2024.07.044

Young, A. M., & Dyer, F. C. (2022). Past experience with spatial or temporal resource unpredictability shapes exploration in honey bees, *Apis mellifera*. Animal Behaviour, 194, 253–264. 10.1016/j.anbehav.2022.09.001

