## Supplementary Materials for "Behavioural flexibility masks delayed costs of environmental instability"

Supplementary Methods

*S1. Relationship between revisitation probability and floral coverage*

The relationship between revisitation probability and the number of unique flowers visited was assessed across individuals and trials. Pearson correlation coefficients were calculated to quantify the association between variables, and partial correlations controlling for total visit number were also computed.

Linear mixed-effects models were fitted to test the effects of revisitation probability and total visit number on the number of unique flowers visited, with bee identity included as a random intercept. Additional models tested the association between revisitation probability and total visit number.

All analyses were conducted using the same dataset and modelling framework as the main analyses.

*S2. Exploratory analyses of pollen extraction and collection*

To explore potential mechanisms underlying treatment effects on pollen collection, we quantified flower-level pollen removal using measurements of pollen mass before and after individual flower visits. Pollen removed from a flower was calculated as the difference between pre-visit and post-visit pollen mass (mg). Because pollen removed from flowers does not necessarily correspond directly to pollen collected by bees, owing to potential losses during handling, grooming, or transport, these analyses were considered exploratory and were conducted to assess whether treatment effects on pollen collection were associated with differences in pollen extraction.

For analyses of the treatment phase (T1–T3), flower-level pollen removal measurements were analysed using linear mixed-effects models including treatment, trial, and their interaction as fixed effects, with colony included as an additional fixed effect and bee identity included as a random intercept. To examine the relationship between pollen extraction and pollen collection, flower-level measurements were aggregated to the bee-trial level by summing pollen removed across all flowers visited by a bee within a trial. Relationships between total pollen removed and pollen load collected by bees were assessed using Pearson correlations and linear mixed-effects models including total pollen removed, treatment, trial, and colony as predictors.

For analyses of the final preference trial (P2), flower-level pollen removal was analysed using linear mixed-effects models with treatment and colony as fixed effects and bee identity as a random intercept. To assess whether persistent treatment differences in pollen collection under identical conditions could be explained by differences in pollen extraction, bee-level summaries of pollen removal were generated by calculating total pollen removed and mean pollen removed per flower for each individual. Associations between pollen removal and pollen load were evaluated using Pearson correlations.

*S3. Orientation-specific carry-over during the post-trial preference test (P2)*

To assess whether persistent treatment differences observed during the post-trial preference test (P2) reflected orientation-specific carry-over from the preceding training phase, we examined whether bees preferentially allocated visits to the flower orientation experienced immediately before P2. For bees in the stable treatment, this corresponded to the orientation presented throughout all three experimental trials (T1–T3), whereas for bees in the unstable treatment it corresponded to the orientation presented during the final training trial (T3).

For each bee, flower visits during P2 were classified as either directed towards the target orientation or towards one of the two alternative orientations. The proportion of visits directed towards the target orientation was analysed using a binomial generalized linear model with treatment and colony included as fixed effects. Because the P2 preference test presented equal numbers of upward-, downward-, and sideways-facing flowers, the null expectation under random visitation was that one-third of visits would be directed towards the target orientation. Estimated marginal means were therefore additionally compared with this chance expectation.

As a complementary analysis retaining orientation-level observations, the number of visits to each flower orientation during P2 was analysed using a negative binomial generalized linear mixed-effects model. Fixed effects included target status (target versus non-target orientation), treatment, their interaction, and colony, with bee identity included as a random intercept. Together, these analyses tested whether the persistent behavioural differences observed during P2 could be explained by preferential revisitation of the previously experienced flower orientation.

Supplementary Results

*S1. Exploratory analyses of pollen extraction and collection*

To investigate potential mechanisms underlying treatment effects on pollen collection, we quantified flower-level pollen removal and examined its relationship with pollen loads collected by bees.

Pollen removal during T1–T3: Patterns of pollen removal changed across the experimental trials, and these temporal changes differed between treatments (Treatment × Trial: $\chi^2_2 = 9.12$, $p = 0.010$). Bees from the stable treatment removed similar amounts of pollen in T1 and T2 but significantly less pollen in T3 (T1–T3: $p = 0.0002$; T2–T3: $p = 0.0002$). Bees from the unstable treatment also exhibited lower pollen removal in T3 than in T1 ($p = 0.034$), although changes across trials were less pronounced. Treatment differences within individual trials were not statistically significant following multiple-comparison correction (all $p > 0.05$).

Across T1–T3, bees that removed more pollen from flowers generally collected larger pollen loads. Total pollen removed per bee-trial was positively correlated with pollen load ($r = 0.60$, $p < 0.001$). After accounting for variation in pollen removal, neither treatment ($\chi^2_1 = 2.20$, $p = 0.138$), trial ($\chi^2_2 = 0.12$, $p = 0.942$), nor the Treatment × Trial interaction ($\chi^2_2 = 1.07$, $p = 0.585$) explained additional variation in pollen load. These results indicate that variation in pollen collection during the treatment phase was closely associated with differences in flower-level pollen removal.

Pollen removal during P2: To assess whether persistent treatment differences in pollen collection under identical conditions could be explained by differences in pollen extraction, we quantified flower-level pollen removal during the final preference trial (P2). Bees from the stable treatment tended to remove more pollen per flower than bees from the unstable treatment (6.93 ± 4.36 mg versus 5.86 ± 4.67 mg, mean ± SD), although this difference was modest and not statistically significant ($\chi^2_1 = 2.72$, $p = 0.099$).

In contrast to the treatment phase, pollen removal and pollen collection were only weakly associated during P2. Mean pollen removed per flower was not significantly correlated with pollen load ($r = 0.23$, $p = 0.228$), and total pollen removed was similarly unrelated to pollen load (r = 0.27, p = 0.157). Thus, although treatment differences in pollen collection persisted under identical conditions, these differences were not strongly explained by variation in flower-level pollen removal.

Taken together, these exploratory analyses suggest that pollen collection during the treatment phase (T1–T3) was closely linked to pollen removal from flowers, whereas the persistent treatment effects observed during P2 were not readily explained by differences in flower-level pollen extraction alone.

*S2. Orientation-specific carry-over during the post-trial preference test (P2)*

To determine whether the persistent treatment differences observed during the post-trial preference test (P2) reflected orientation-specific carry-over from the preceding training phase, we tested whether bees preferentially allocated visits to the orientation experienced immediately before P2. For bees in the stable treatment this corresponded to the orientation experienced throughout T1–T3, whereas for bees in the unstable treatment it corresponded to the final orientation experienced during T3.

There was no evidence that the proportion of visits directed towards the previously experienced orientation differed between treatments (Treatment: $\chi^2_1 = 0.017$, $p = 0.897$; Fig. S4). Estimated probabilities of visiting the target orientation were similar for bees from the stable (0.281, 95% CI: 0.20–0.38) and unstable treatments (0.276, 95% CI: 0.19–0.39). Furthermore, neither treatment showed evidence that visits to the previously experienced orientation exceeded the expectation under random visitation (one-third of visits; stable: $p = 0.287$; unstable: $p = 0.303$).

A complementary negative binomial mixed-effects model analysing orientation-level visit counts likewise found no evidence that bees preferentially visited the previously experienced orientation within either treatment (target status: $p = 0.629$ for stable bees; $p = 0.899$ for unstable bees). Instead, visit allocation during P2 was distributed similarly across flower orientations irrespective of previous experience.

Together, these analyses indicate that the persistent treatment differences observed during P2 were not explained by preferential revisitation of the previously experienced flower orientation, suggesting that the carry-over effects arose from broader changes in pollen-foraging behaviour rather than orientation-specific visitation preferences.

Supplementary Tables

**Table S1. Distribution of flower orientations across trials (T1–T3) in the unstable treatment**

| **Orientation** | **T1** | **T2** | **T3** | **Total** |
|---|---|---|---|---|
| Downward | 4 | 5 | 7 | 16 |
| Sideways | 7 | 6 | 3 | 16 |
| Upward | 5 | 5 | 6 | 16 |
| **Total** | 16 | 16 | 16 | 48 |

Number of bees exposed to each flower orientation in each trial (T1–T3) within the unstable treatment (n = 16 bees). Each bee experienced all three orientations once. The distribution of orientations across trials was approximately balanced across individuals.

**Table S2. Results of morphology-augmented models testing the effects of bee body mass and intertegular span (ITS) on behavioural and performance metrics.**

| Response variable | Predictor | $\chi^2$ / F | df | p-value |
|---|---|---|---|---|
| Revisitation | Body mass | 0.0017 | 1 | 0.967 |
| | ITS | 0.1455 | 1 | 0.703 |
| Pollen load | Body mass | 0.2183 | 1 | 0.640 |
| | ITS | 0.4063 | 1 | 0.524 |
| Bout duration | Body mass | 0.5969 | 1 | 0.440 |
| | ITS | 2.8628 | 1 | 0.091 |
| Foraging efficiency | Body mass | 0.0959 | 1 | 0.757 |
| | ITS | 0.0876 | 1 | 0.767 |
| Sonication probability | Body mass | 3.9977 | 1 | 0.046* |
| | ITS | 0.7002 | 1 | 0.403 |

Morphology-augmented versions of the main models included bee body mass and intertegular span (ITS) as fixed effects. Values shown are Type III Wald chi-square statistics (generalized linear mixed models) or equivalent tests for linear mixed models. Body mass showed a weak effect on sonication probability, but neither morphological trait significantly predicted revisitation behaviour, pollen load, bout duration, or foraging efficiency.

Supplementary Figures

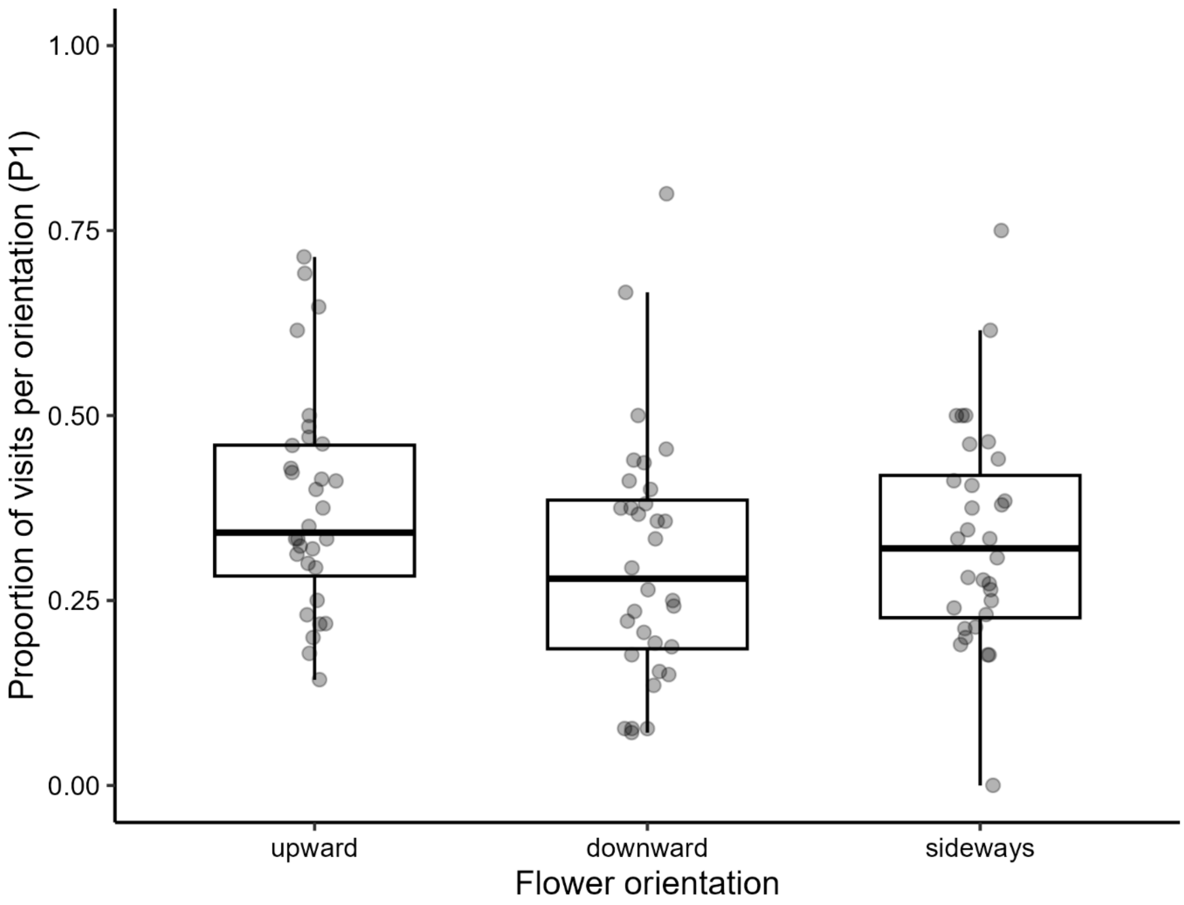


**Figure S1. Baseline orientation preference in the retained subset (P1).** Proportion of visits to flowers of different orientations (upward, downward, sideways) during the initial preference test (P1) for individuals retained in the main experiment (n = 32). Points represent individual-level proportions, and boxplots show medians and interquartile ranges. There was no strong evidence that visitation differed across orientations (GLMM: $\chi^2_2 = 5.66$, $p = 0.059$; Friedman test: $\chi^2_2 = 2.09$, $p = 0.352$), indicating no clear baseline preference prior to experimental manipulation.

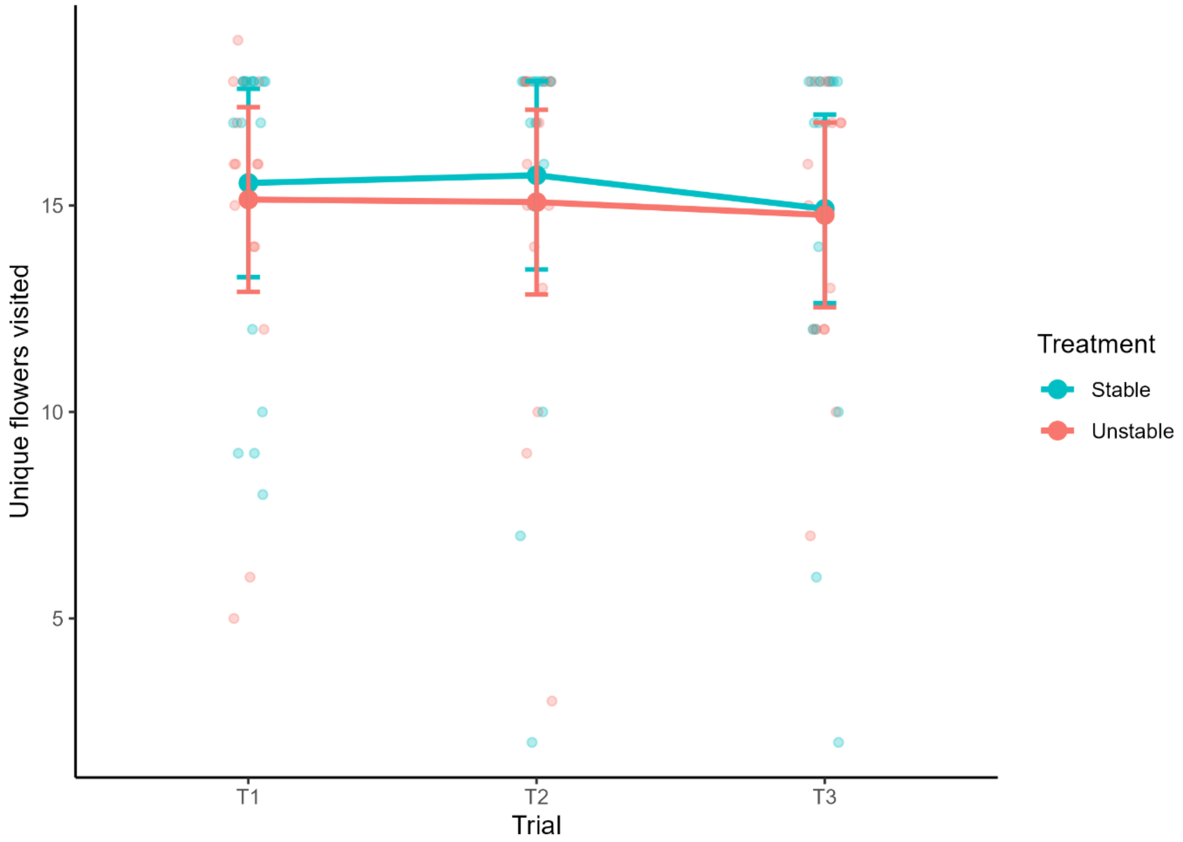


**Figure S2. Floral coverage across trials and treatments.** Number of unique flowers visited per bout (floral coverage) across three consecutive trials (T1–T3) under stable and unstable conditions. Points represent individual observations, with slight horizontal jitter for clarity. Larger points and error bars indicate model-estimated marginal means ± 95% confidence intervals. There was no evidence that floral coverage differed between treatments or across trials, indicating that exploration of floral resources remained stable despite environmental manipulation.

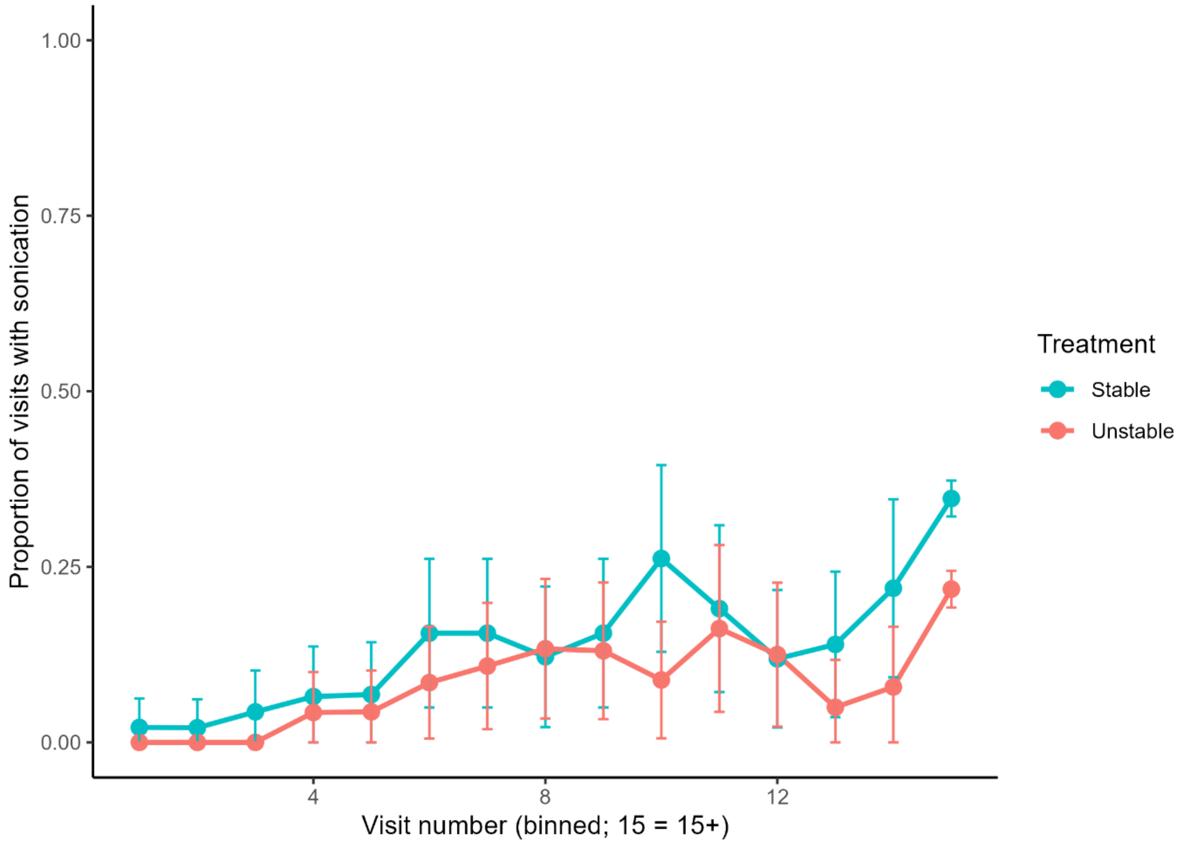


**Figure S3. Within-bout dynamics of sonication behaviour.** Proportion of visits involving sonication as a function of visit number within a foraging bout (binned; 15 = 15+), shown separately for stable and unstable treatments. Points represent model-estimated means ± 95% confidence intervals. Sonication probability increased nonlinearly across visits within bouts and differed between treatments, consistent with experience-dependent modulation of handling behaviour during foraging sequences.

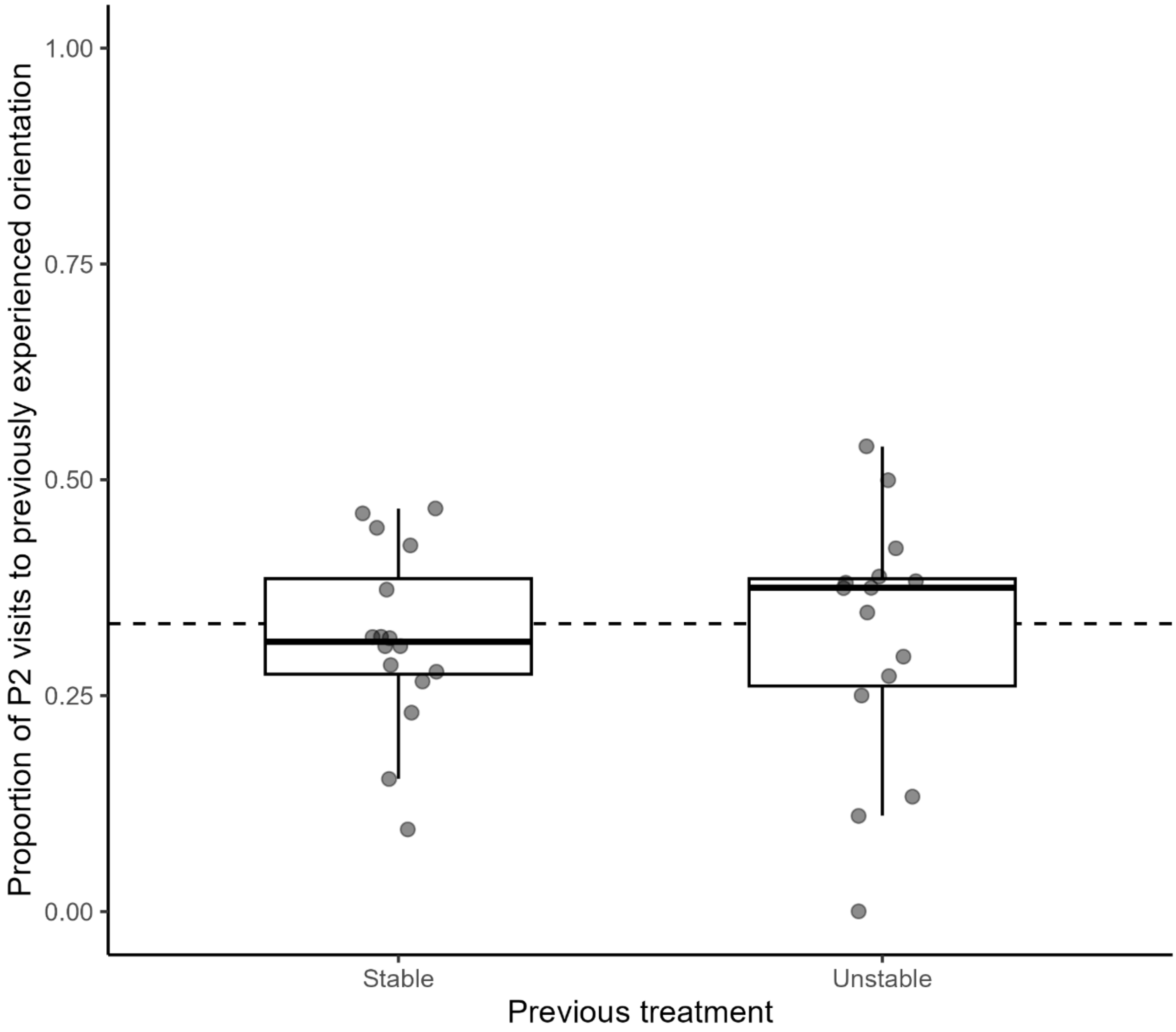


**Figure S4. Orientation-specific carry-over during the post-trial preference test (P2).** Proportion of flower visits directed towards the previously experienced flower orientation during P2. For bees in the stable treatment, the previously experienced orientation corresponded to the orientation presented throughout the three experimental trials (T1–T3), whereas for bees in the unstable treatment it corresponded to the orientation experienced during the final training trial (T3). Points represent individual bees, and boxplots show medians and interquartile ranges. The dashed line indicates the expected proportion of visits under random visitation (1/3). There was no evidence that bees preferentially visited the previously experienced orientation or that orientation-specific visitation differed between treatments.
